# Ecological dynamics of the Atlantic salmon gut microbiota across developmental phases and geographic regions

**DOI:** 10.1101/2025.09.30.679454

**Authors:** Wasimuddin, Håkon Pedersen Kaspersen, Snorre Gulla, Pimlapas Leekitcharoenphon, Frederik Duus Møller, Samantha White, Simon MacKenzie, Arne Holst-Jensen, Ottavia Benedicenti

**Affiliations:** Food Safety and Animal Health Research, Norwegian Veterinary Institute, Ås, Norway; Fish Health Research, Norwegian Veterinary Institute, Ås, Norway; Research Group for Genomic Epidemiology, National Food Institute, Technical University of Denmark, Lyngby, Denmark; Marine Institute, Rinville, Co. Galway H91 R673, Ireland; Institute of Aquaculture, University of Stirling, Stirling FK9 4LA, UK

**Author notes:** **Correspondence** Wasimuddin.

**Keywords:** Atlantic Salmon, 16S rRNA gene, developmental phases, geographic regions, microbial diversity, community composition

## Abstract

The gut microbiota is vital to host health, yet the relative influence of host traits and environmental factors on fish gut microbiota dynamics remains underexplored. We investigated the ecological dynamics of Atlantic salmon (*Salmo salar*) gut microbiota, by analysing 847 samples from wild and farmed salmon across diverse geographic regions, developmental phases, and associated diet and environmental microbiota. Farmed salmon exhibits reduced microbial diversity and distinct community composition with increased Firmicutes and reduced Proteobacteria compared to wild salmon. Microbial diversity declined with advancing developmental phases notably due to reduced Proteobacteria and expanded *Mycoplasma*. Diet was the primary contributor (∼23%) to farmed salmon microbiota, with environmental inputs varying by region and phase. These findings highlight the importance of aquaculture practices guided by microbiota insights, while emphasize the need to preserve microbial diversity in wild populations to enhance resilience against environmental pressures, contributing to both sustainable farming and conservation strategies.

## Introduction

The gut microbiota plays a pivotal role in host health, influencing diverse physiological processes such as nutrient absorption, immune system development, disease resistance, and overall homeostasis^1^. In fish, these microbial communities are highly dynamic, shaped by complex interactions with environmental factors, dietary inputs, and intrinsic host factors, including developmental phase and genetic background^2,3^. Understanding the diversity and structure of the gut microbiota is particularly critical in aquaculture, where promoting optimal gut health is essential for enhancing productivity, bolstering disease resilience, and ensuring sustainability^4,5^. Moreover, enhanced knowledge of gut microbiota in marine fishes is vital for conserving wild fish populations, which face unique ecological challenges and environmental pressures^6^.

Despite an increasing interest in the fish microbiota, our understanding of the relative importance of the effects of host traits versus environmental factors on shaping fish microbiota is still largely unknown^7,8^. Such knowledge is essential though, as changes that cause disturbances in the composition or abundance pattern of microbial communities beyond the natural range might cause dysbiosis affecting the health and growth of fish^9^. Environmental factors such as habitat type, diet, and salinity play important roles in shaping fish gut microbiota, since aquatic environment subjects fish to continuous, direct exposure to waterborne microbial communities^8,10^. Farmed fish live in controlled environments with standardized diets; consequently, it can be hypothesized that their gut microbiota differ from those of wild fish, which are exposed to diverse natural microbial pools and varied diets^11^. Similarly, host intrinsic factors such as developmental phase transitions and genetics could also influence the gut microbiota of fish^7^. Specifically, little is known about how developmental phase associated physiological transitions, from early development phases to late growth phase, shape the gut microbiota in fishes at varying geographical locations. Thus, to elucidate the baseline inter-individual variability in fish gut microbiota from undesirable shifts in microbiota diversity and compositions, we need to account simultaneously for intrinsic host traits and extrinsic environmental factors^7,10^.

The Atlantic salmon (*Salmo salar*) is one of the most economically significant aquaculture species and exhibits a complex life cycle marked by distinct physiological transformations. Under natural conditions, these anadromous fishes hatch in freshwater rivers, where they may remain for up to five years before undergoing smoltification, a process that prepares them for migration to the ocean and subsequent maturation. The transition from freshwater to seawater presents new environmental and dietary challenges. They typically spend one to four years in the marine environment before returning to their natal rivers to spawn^12^. These physiological transitions, coupled with interactions with diet and environmental exposure, make Atlantic salmon an ideal model for studying gut microbiota dynamics. Due to its long distance migrations between freshwater and ocean habitats, wild Atlantic salmon populations face multiple threats such as habitat degradation, climate change, and interactions with escaped farmed salmon^12^. Individual salmon return to spawn in their natal river, which has enabled the formation of genetically distinct populations, adapted to the local conditions among and within river systems^13^. As the gut microbiota contributes to environmental adaptation and microbial diversity is critical for the resilience of wild populations^14^, gaining insight into the gut microbiota of wild salmon is of paramount importance. Farmed Atlantic salmon hatch and develop through smoltification in freshwater facilities, before sea transfer and on growth in large pens, with slaughtering taking place about three years after hatching. Farmed salmon are subject to selective breeding over multiple generations, limited environmental variability, and standardized diets optimized for rapid growth and high yield^15^. While enhancing productivity, these conditions can lead to distinct gut microbiota profiles^16^, which can further influence immune function and disease resilience^17^. Indeed, farmed salmon in seawater environments are particularly susceptible to parasitic infestations, such as sea lice, which may be transmitted to wild salmonid populations through waterborne dispersal, direct contact, or escape events^18^. The limited environmental exposure and standardized diets in salmon farming systems provide an opportunity to assess the relative contributions of diet and environment in shaping the gut microbiota of farmed salmon. Gaining such insights is essential, as understanding microbiota dynamics in both farmed and wild salmon underpins the development of sustainable aquaculture practices and effective conservation strategies.

To address these knowledge gaps, this study aims to investigate: (i) How microbial diversity, community composition and taxa abundance differ between wild and farmed salmon, and how these differences are influenced by geographic origin; (ii) The effects of developmental phases on gut microbiota diversity and composition in farmed salmon, including the identification of developmental phase-specific taxa; (iii) Relative contributions of diet and environmental factors (e.g., freshwater, seawater) to the gut microbiota of farmed salmon, considering geographic and developmental phase variation. We explore these questions by examining the gut microbiota of wild and farmed Atlantic salmon (*Salmo salar*) across multiple developmental phases and geographic regions (Norway, Ireland, Scotland), along with diet and environment (freshwater, seawater) samples. This comprehensive analysis of the ecological dynamics of the salmon gut microbiota provides actionable insights for improving sustainable aquaculture practices and informing conservation strategies for wild salmon populations.

## Materials and Methods

### Sample collection

Samples of wild and farmed Atlantic salmon, as well as diet and environment (i.e. freshwater, seawater) of farmed salmon were collected from Ireland, Scotland and Norway. Specifically, 273 samples from Ireland, 379 from Norway and 195 from Scotland (total=847) were retained after all laboratory and bioinformatic procedures, representing diet (diet=33), environment (freshwater=30, seawater=35), and fish (early developmental phases=31, digesta=304, intestinal tissue=414). Based on their biological significance during the Atlantic salmon life cycle, nine sampling timepoints were defined: fertilised eggs (T0), yolk sac larvae (T1), fry (T2), parr/pre-vaccination (T3), post-vaccination (T4), pre-smoltification (T5), post-smoltification (T6), pre-seawater transfer (T7), post-seawater transfer (T8), and late growth phase (T9). All timepoints were included in our study except fry (T2), pre-smoltification (T5), and pre-seawater transfer (T7), owing to discrepancies in these sample collections across countries. For the early developmental phases of farmed salmon (SDP), i.e., fertilised eggs and yolk sac larvae, for each sampling 100 fertilized eggs (ca. 300/400-degree days after fertilization) or 100 yolk sac larvae (ca. 700/800-degree days after fertilization) were randomly collected and transferred to four sterile stomacher bags of 25 eggs/larvae per bag, each corresponding to one sample. The bag content was then homogenized in a stomacher (Seward Stomacher^®^ 80) at normal speed for 1 min, and the homogenate was stored in a sterile plastic tube at -80ºC until DNA extraction. For all later SDPs, individual fish were randomly sampled. Tricaine methanesulfonate (MS-222, Sigma-Aldrich, USA) was used to euthanize (0.3-0.5 g/L) Atlantic salmon for sampling. The wild Atlantic salmon were caught from each country and similarly sampled after euthanisation. The entire gastrointestinal tract was dissected and cut into three sections; anterior, mid and posterior from each individual fish. Digesta (25 g) from the anterior and mid intestines was separated from the tissue. Empty intestine segments were washed 3-4 times in sterile phosphate-buffered saline (PBS, Sigma-Aldrich, USA). The intestine and digesta samples were then stored in sterile tubes at - 80ºC until DNA extraction.

Environment sampling (i.e. water) was conducted before other operational activities (e.g., fish sampling) to minimize transient increases in contamination. Samples were collected from either the freshwater tank or the seawater cage. One litre of water was taken for each sample and pre-filtered using a 20 µM filter to remove larger sized particles before final filtering through a Sterivex^TM^ filter (pore-size 0.22 µM, Millipore, USA) using a peristaltic pump. The Sterivex^TM^ filter was stored in a sterile plastic bag at -80ºC until DNA extraction. For the diet samples, five random feed samples of 10 g each were collected with a sterile scoop at the farm. The feed samples were mixed with 100 mL peptone water and were homogenized in the stomacher for 1 min at normal speed, filtered (same procedure as the water filtering), and 25 mL of the homogenate was stored in a sterile plastic tube at -80ºC.

### DNA Extraction and 16S rRNA Gene Amplicon Sequencing

Total DNA was isolated using the DNeasy PowerSoil Pro Kit (Qiagen, USA). Samples were processed by 16S rRNA gene V3-V4 region amplification through PCR followed by sequencing using Illumina’s recommended protocol (https://support.illumina.com/). A two-round amplification process was used to amplify the DNA samples, while reducing dimer formation, which is often a problem in multiprimer, multitemplate PCR, especially with primers containing long overhang regions^19^. Briefly, a PCR was performed with the use of the primers: 341F-5’TCGTCGGCAGCGTCAGATGTGTATAAGAGACAGCCTACGGGNGGCWGCAG, 805R-5’GTCTCGTGGGCTCGGAGATGTGTATAAGAGACAGGACTACHVGGGTATCTAATCC, carrying overhang adapters for compatibility with Illumina index and sequencing adapters^20,21^. PCR reactions were run in 25 µL reactions using KAPA HiFi Hotstart (KAPA Biosystems, Roche, USA), 1 µM of each primer and 5 ng/µL DNA template, using the following thermal profile: 95°C for 3 min; 25 cycles of 95°C for 30 s, 55°C for 30 s, 72°C for 30 s; and 72°C for 5 min. The second PCR Index reaction was run in a 25 μL reaction with KAPA Hifi HotStart and Illumina Nextera indexing primers (95°C for 3 min; 8 cycles of 95°C for 30 s, 55°C for 30 s, 72°C for 30 s; and 72°C for 5 min). Both rounds of amplicon libraries were purified with Beckman Coulter™ Agencourt AMPure XP (Thermo Fisher Scientific, USA) before proceeding to the next analysis step. Barcoded purified products were validated on Agilent Bioanalyzer and quantified using Nanodrop (Thermo Fisher Scientific, USA). The final pooled libraries at concentration of 4 nM in hybridization buffer were paired-end sequenced (2 x 300 cycles) using MiSeq Reagent Kit-v3 on a MiSeq Illumina instrument at the Denmark Technical University (DTU) sequencing facility. Sequence data were deposited in the European Nucleotide Archive under project ID PRJEB81272.

### Bioinformatic analysis

Demultiplexed reads without barcodes and adapters were received as output from the Illumina sequencing platform. For data pre-processing, we employed a nf-core^22^ NextFlow pipeline: AmpliSeq (version 2.4.1)^23^. In brief, reads were trimmed from both ends based on the quality profile; error rates were obtained from the data by using the parametric error model as implemented in DADA2^24^. After the reads had been denoised and merged, chimeric sequences were removed from the dataset by following the “consensus” method. Thus, non-chimeric amplicon sequence variants (ASVs; i.e., sequences differing by as little as one nucleotide) were identified in each sample. The taxonomy of representative ASVs was assigned using the naïve Bayesian classifier method with the SILVA v138 non-redundant database. All subsequent analyses were performed within the R environment (version 3.5.3)^25^. The Phyloseq (version 1.26.1)^26^ package was used for further data processing, with ASVs belonging to chloroplast, mitochondria, Eukaryota, and unassigned ASVs at the phylum level being removed from the dataset. We further removed the samples with less than 1,000 reads as well as ASVs showing less than 10 reads from the overall dataset. We recovered, on average, 43,356 (range 1,015–901,417) high-quality reads per sample after all processing steps.

## Statistical analysis

### Alpha diversity analysis

To determine inter-individual diversity, we calculated two alpha diversity measures: observed number of ASVs (Observed ASVs) and Shannon diversity using the Phyloseq package. Samples collected from different locations (anterior, mid or posterior within each compartment–digesta or intestinal tissue), were categorized broadly as digesta and tissue samples to reduce variability, improve statistical power, and focus on broad-scale differences related to salmon domestication status (SDS) (wild versus farmed) and developmental phases (SDPs). To successively account for the potential effect of known variables on diversity measures and achieve a balanced design, we constructed three different generalized linear mixed models (GLMMs). In the first model, we accounted for the difference in wild versus farmed salmon microbiota in their alpha diversity indices by modelling alpha diversity according to SDS (Wild, n □= □169; Farmed, n □= □289), country (Ireland, n □= □109; Norway, n □= □240; Scotland, n □= □109), sample category (Digesta, n = 179; Intestinal tissue, n = 279) and interaction between these (SDS*country*sample category). We also added timepoint (T6, n □= □158; T9, n □= □300) in the model, mainly to control its potential effect. In the second model for farmed salmon, we tested the effect of timepoints (SDP) on alpha diversity indices by utilizing similar timepoint–collected samples of salmon across countries. For this we included timepoint (T0, n = 15; T1, n = 16; T3, n = 89; T4, n = 73; T8, n = 98; T9, n = 248), country (Ireland, n □= □181; Norway, n □= □278; Scotland, n □= □80), and interaction between these (timepoint*country). In the third model, we compared microbial alpha diversity indices among diet, environment and salmon gut (i.e. digesta, intestinal tissue) for farmed salmon. For this model, we included sample category (Diet, n = 33; Freshwater, n = 30; Seawater, n = 35; Digesta, n = 229; Intestinal tissue, n = 279), country (Ireland, n □= □201; Norway, n □= □295; Scotland, n □= □110), and interaction between these (sample category*country). We also added timepoint (T3, n □= □112; T4, n □= □95; T8, n □= □120; T9, n □= □279) in the model, mainly to control its potential effect. In all three GLMMs, we added sequencing depth to account for differential sequencing effort between samples and sequencing run to control the random effect of batch processing of samples^27^. To facilitate model convergence, sequencing depth was scaled. We used a log distribution for modelling the count data (Observed ASVs) and a normal distribution for modelling the continuous data (Shannon diversity). Model selection was based on the information-theoretic (IT) approach using a second-order Akaike’s Information Criterion corrected for small sample sizes (AIC_C_) as an information criterion and Akaike weights (ω) to determine model support^28^. For all GLMMs, we report both conditional and marginal coefficients of determination of each model: R^2^_GLMM(c)_, which explains the variance of both the fixed and random factors, and R^2^_GLMM(m)_, which explains the variance of the fixed factors only. The latter was calculated as the variance explained by the best model and the ΔAIC_C_. The conditional parameter estimates (*β*) and confidence intervals (95% CI) were calculated, and we report back-transformed values in cases where log transformation was employed.

### Beta diversity analysis

To assess the bacterial community composition between samples, we calculated the Bray– Curtis distance matrices using Phyloseq. Bray–Curtis accounts for the presence-absence of the taxa and gives weight to taxa abundance. We tested for differences in microbial beta diversity among categories using the permutational multivariate analysis of variance (PERMANOVA) test with 999 permutations implemented in the adonis function of the vegan package (version 2.6.2)^29^. We built three statistical models with variables as for alpha diversity and retained sequencing depth and sequencing run variable in our full models to statistically account for their model support. To visualize patterns of separation between different sample categories, Principal Coordinates Analysis (PCoA) plots were prepared based on the Bray–Curtis distance.

### Differential abundance analysis for major phyla and ASVs

In order to evaluate differences in the relative abundance of the five major bacterial phyla (>0.1%) between farmed and wild salmon across countries, we applied similar GLMMs as done for alpha diversity. Briefly, we ran the GLMM model for each phylum abundance by accounting SDS (Wild, n □= □169; Farmed, n □= □289), country (Ireland, n □= □109; Norway, n □= □240; Scotland, n □= □109), sample category (Digesta, n = 179; Intestinal tissue, n = 279), interaction between these (SDS*country*sample category) and timepoint (T6, n □= □158; T9, n □= □300). Similarly, to understand the effect of timepoints (SDPs) on the abundance of major bacterial phyla in farmed salmon, we applied the GLMM for each of the five major phyla, accounting for timepoint (T0, n = 15; T1, n = 16; T3, n = 89; T4, n = 73; T8, n = 98; T9, n = 248), country (Ireland, n □= □181; Norway, n □= □278; Scotland, n □= □80), and their interaction (timepoint*country). In all the above models, we added sequencing depth (scaled) to account for differential sequencing effort between samples and sequencing run to control the random effect of batch processing of samples as in alpha diversity models^27^. Model selection was based on the IT approach using a second-order AIC_C_ as an information criterion and Akaike weights (ω) to determine model support^28^. The conditional parameter estimates (*β*) were calculated, and we report back-transformed values in cases where log transformation was employed.

In order to identify the ASVs accountable for differences in the microbiota of wild and farmed salmon across countries, we employed a negative binomial-model-based approach; available in the *DESeq2* package^30^ after removing spurious ASVs (i.e., ASVs present in less than three samples). Wald tests were performed and only ASVs remaining significant (p≤0.01) after the Benjamini–Hochberg correction were retained^31^.

To detect ASVs that exhibit linear trends of increase, decrease or mid-peak during the salmon SDPs, we employed ANCOM-BC2^32^. ANCOM-BC2 was specifically developed to detect trends in taxa abundance based on linear mixed effect models with bias correction through the default ‘ancombc2’ function. To have robust estimates and to detect major trends in ASVs abundance, we merged the timepoint data into major life-stages: T0-T1 as ‘early’, T5-T6 as ‘mid’ and T8-T9 as ‘late’. We accounted for country and SDP and kept reads per sample as a fixed variable and controlled the random effect of sequencing batches in the model. Significant ASVs (p≤0.05) after Holm–Bonferroni correction were reported^33^.

To understand the relative contribution of the dietary and environmental (freshwater or seawater) microbiota in shaping the salmon gut microbiota (digesta or intestinal tissue), we ran a Bayesian analysis approach implemented in SourceTracker with default parameters, together with 100 burnings^34^. SourceTracker quantitatively assigns proportions of the microbial contaminants in a digesta/intestinal tissue microbiota to a set of potential sources represented by the source microbiota (diet, freshwater or seawater). The part of the salmon gut microbiota that could not be assigned to any of the potential sources was collectively classified as from ‘unknown’ sources.

## Results

### Microbiota diversity and community composition differ between wild and farmed salmon

To test whether the salmon intestinal microbiota (digesta or intestinal tissue) differs between wild and farmed fish (Salmon Domestication Status; SDS), we calculated the microbial alpha diversity indices, specifically Observed ASVs and Shannon diversity for all individuals. Using model selection based on the IT approach, we found strong support for an effect of SDS, country and sample category, but also a combined effect of these three (SDS*country*sample category) and timepoint on Observed ASVs (ΔAICc = □274.13, R^2^_GLMM(m)_ = □0.499, R^2^_GLMM(c)_ = □0.763, Figure 1a, Supplementary Figure 1a). Similarly, Shannon diversity was noted to be influenced by SDS, country, sample category, timepoint and interaction terms (country*SDS, country*sample category) but not by the combined effect of SDS*country*sample category (ΔAICc = □177.67, R^2^_GLMM(m)_ = □0.484, R^2^_GLMM(c)_ = □0.601, Figure 1b, Supplementary Figure 1b). For both alpha diversity indices, in general, lower diversity was observed in the farmed as compared to wild salmon (Observed ASVs: *β*_*W*_ □= □214.5 (95% CI □= □31.544–1441.986), *β*_*F*_ □= □155.8 (95% CI □= □64.615–370.482); Shannon: *β*_*W*_ □= □3.7 (95% CI □= □1.369–5.234), *β*_*F*_ □= 2.9 (95% CI □= □2.086–3.875)). Salmon from Norway showed the highest diversity, followed by Ireland and Scotland (Observed ASVs: *β*_*N*_ □= □575.9 (95% CI □= □ 44.655–3645.06), *β*_*I*_ □= □155.8 (95% CI □= □64.615–370.482), *β*_*S*_ □= □109.0 (95% CI □= □16.104–732.999); Shannon: *β*_*N*_ □= □ 4.3 (95% CI □= □2.511–6.204), *β*_*I*_ □= 2.9 (95% CI □= □2.086–3.875), *β*_*S*_ □= 2.5 (95% CI □= □ 0.643–4.471)). Digesta samples showed a higher diversity than intestinal tissue samples (Observed ASVs: *β*_*In*_ □= □140.7 (95% CI □= □41.138–475.087), *β*_*D*_ □= □155.8 (95% CI □= □64.615–370.482); Shannon: *β*_*In*_ □= □2.5 (95% CI □= □1.063–3.969), *β*_*D*_ □= 2.9 (95% CI □= □2.086–3.875)) (Figure 1a, b). Overall, the results demonstrated a strong effect of SDS on microbial alpha diversity indices, although this effect was comparatively less pronounced when considering ASV abundance (i.e., Shannon).

**Figure 1.**
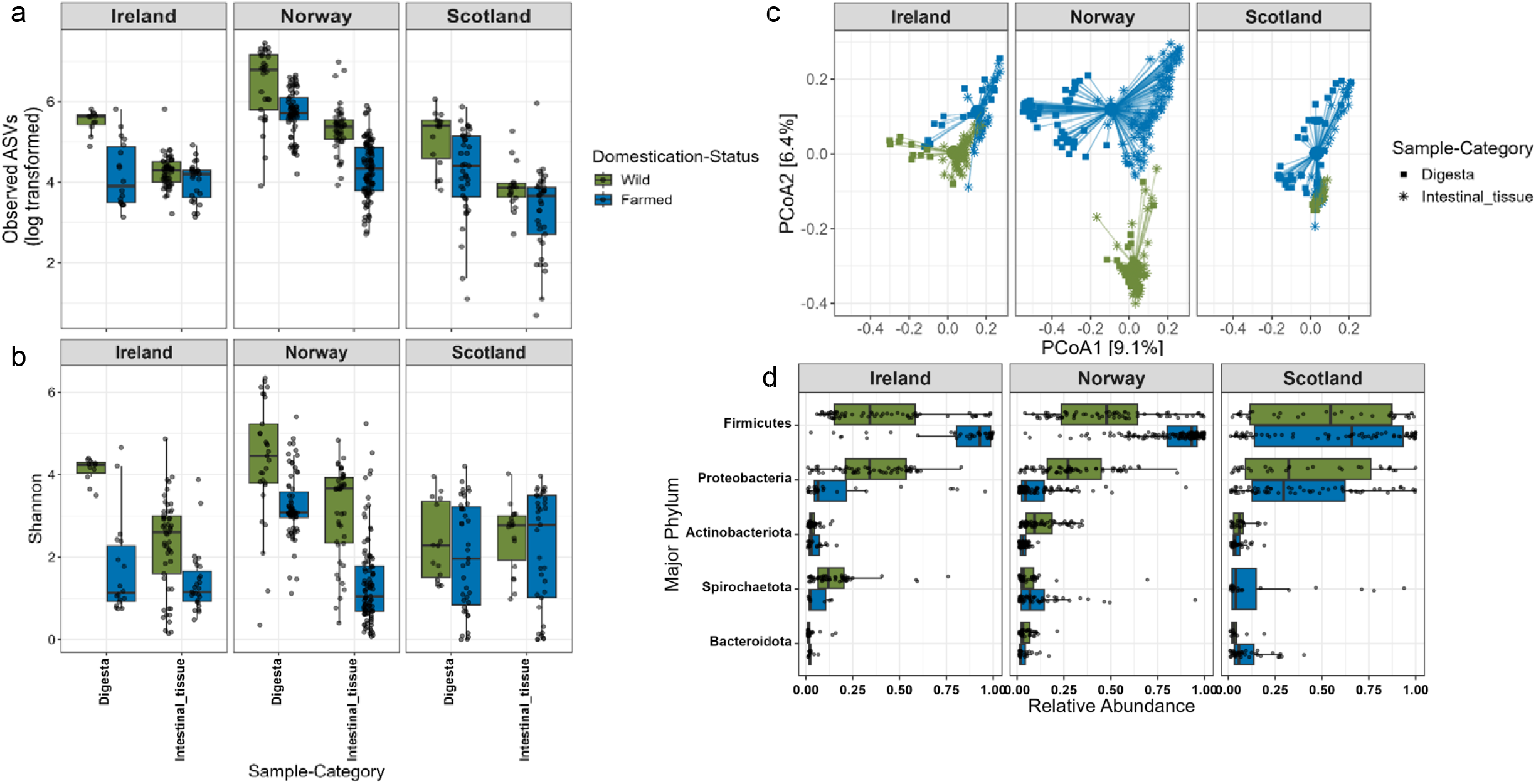
Wild and farmed salmon gut microbiota differ in diversity and composition across countries. (**a**) Box plots showing Observed ASV (natural log transformed) and (**b**) Shannon diversity in digesta and intestinal tissue samples for wild and farmed salmon across countries. The jittered dots represent individual values of samples. (**c**) The PCoA plot reflecting microbial community composition of digesta and intestinal tissue of wild and farmed salmon across country, with sample centroids per group. Percentages of explained variance by each principal axis are indicated in square brackets. (**d**) Box plot showing three major phyla (>0.1%), specifically Firmicutes (ΔAIC_C_ = 35·11), Proteobacteria (ΔAIC_C_ = 18·04) and Bacteroidota (ΔAIC_C_ = 12·61) differ in relative abundance between wild and farmed salmon. The jittered dots represent individual values of samples.

We also assessed the influence of SDS on microbial community composition. While sequencing run (R^2^ = □0.162, p = □0.001) and sequencing depth (R^2^ = □0.004, p = □0.001) did impact the community compositions, the PERMANOVA models indicated a strong support for the influence on the gut microbial beta diversity estimates by: SDS (R^2^ = □0.089, p = □0.007), country (R^2^ = □0.071, p = □0.001), sample category (R^2^ = □0.053, p = □0.001), and the interaction between them (SDS*country*sample category) (R^2^ = 0.020, p = □0.001) (Figure 1c). Additionally, timepoint (R^2^ = □0.022, p = □0.001) and other interactions (country*SDS; R^2^ = □0.002, p = □0.001, SDS*sample category; R^2^ = □0.012, p = □0.001) also influenced the community composition. In the PCoA plots across countries, 15.5% variation in PCoA axes could be explained by SDS and sample category (Figure 1c). Furthermore, wild salmon from different countries showed differential spread in their gut microbiota composition in the PCoA plot (Supplementary Figure 2).

### Differentially abundant major phyla and ASVs between wild and farmed salmon

The SDS (i.e. wild, farmed) influenced the relative abundance of taxa at various taxonomic levels. At the higher phylum level (>0.1%), three phyla, namely, Firmicutes (ΔAIC_C_ = □35.11), Proteobacteria (ΔAIC_C_ = □18.04), and Bacteroidota (ΔAIC_C_ = □12.61), were detected as being differentially abundant between wild and farmed salmon after controlling for sequencing depth and sequencing run in the applied GLMMs (Figure 1d). Among these observed phyla, Firmicutes and Bacteroidota increased whereas Proteobacteria decreased in relative abundance in farmed salmon in comparison to wild salmon (Firmicutes: *β*_*W*_ □= □0.52, *β*_*F*_ □= □0.74; Bacteroidota: *β*_*W*_ □= □0.01, *β*_*F*_ □= □0.02; Proteobacteria: *β*_*W*_ □= □0.23, *β*_*F*_ □= □0.06) (Figure 1d).

At ASV level, by applying negative binomial model-based approaches, we identified 71 (Farmed: decrease=58, increase=13), 135 (Farmed: decrease=63, increase=72) and 103 (Farmed: decrease=40, increase=63) ASVs differentially abundant (p≤0.01) between wild and farmed salmon for Ireland, Norway and Scotland, respectively (Figure 2a). We found 21 ASVs which are common in wild-farmed salmon comparisons across three countries. Among these 21, three ASVs belonging to genus *Mycoplasma, Aliivibrio* and *Photobacterium* showed increased abundance in farmed salmon, whereas two ASVs belonging to genus *Brevinema* and *Rickettsia* showed decreased abundance in farmed salmon in all three countries (Figure 2b).

**Figure 2.**
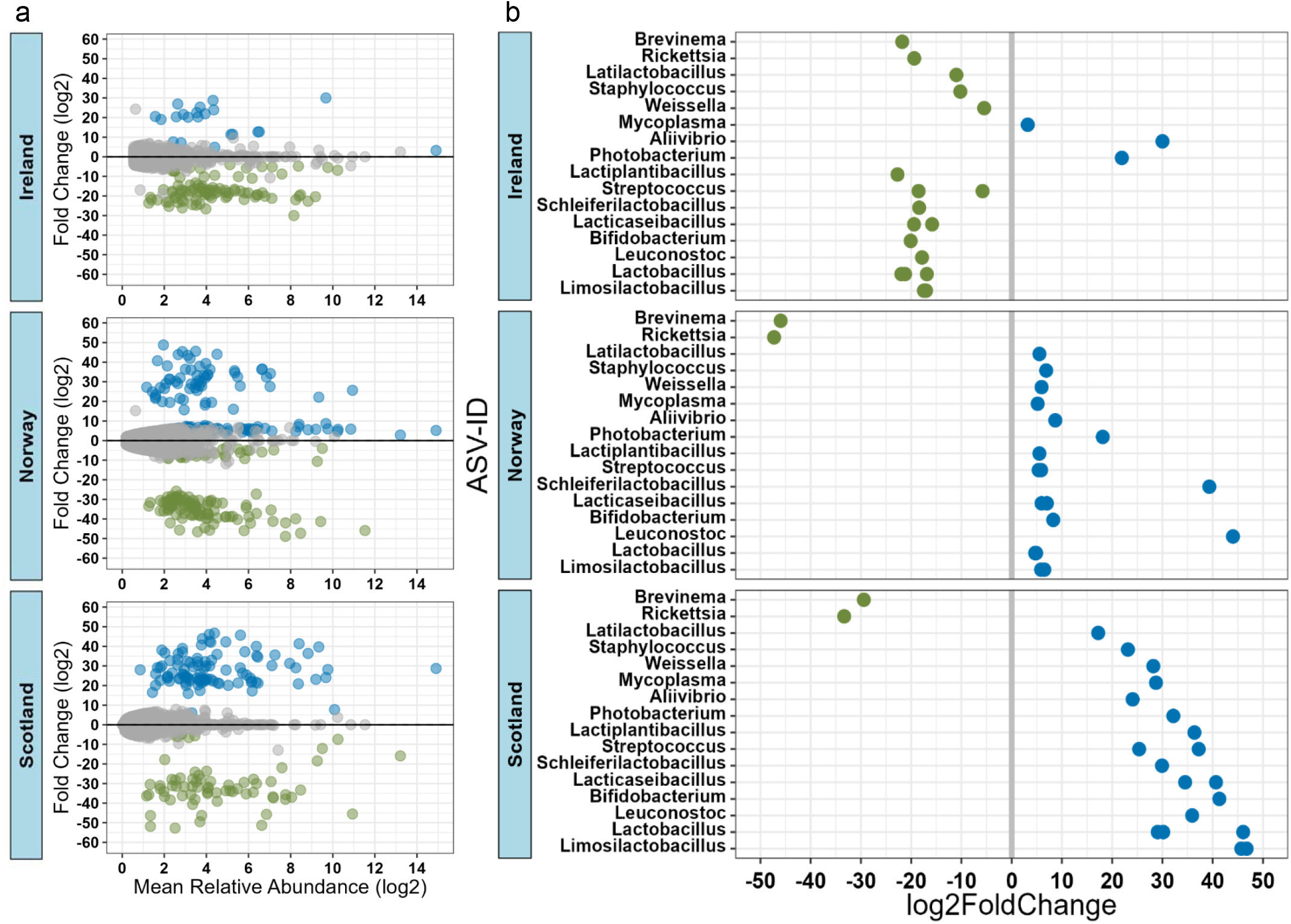
Differentially abundant ASVs between wild and farmed salmon gut microbiota across countries. (**a**) The MA plots display the log2-fold change of all ASVs and their log-mean abundance plotted on y- and x-axes, respectively, for each country. ASVs being differentially abundant between wild and farmed were determined by DESeq2 analysis (Benjamini–Hochberg correction, p≤0.01). Colours refer to enriched ASVs in wild (green) or farmed (blue) salmon for all comparisons (grey=nonsignificant). (**b**) Common ASVs (n=21) across countries for wild and farmed comparison are shown with their respective genus identity at y-axes and log2-fold change on x-axes. Colours refer to enriched ASVs in wild (green) or farmed (blue) salmon for all comparisons.

### Salmon developmental phases influence microbiota diversity and community composition

The GLMM analyses to assess the effect of timepoint (Salmon Developmental Phases; SDP) on farmed salmon across three countries, yielded strong support for timepoint and country, but also combined effect (timepoint*country), of these on Observed ASVs (ΔAICc = □36.38, R^2^_GLMM(m)_ = □0.432, R^2^_GLMM(c)_ = □0.500) as well as on Shannon (ΔAICc = □25.80, R^2^_GLMM(m)_ = □0.406, R^2^_GLMM(c)_ = □0.489) (Figure 3a, Supplementary Figure 3). Overall, gut bacterial diversity showed a decreasing trend in successive SDPs of farmed salmon, regardless of variances in gut microbial diversity at different SDPs across countries (Observed ASVs: *β*_*T0*_ □= □ 850.8 (95% CI □= □ 261.886–1243.831), *β*_*T1*_ □= □202.5 (95% CI □= □ 21.623–852.0), *β*_*T3*_ □= □144.9 (95% CI □= □20.381–464.865), *β*_*T4*_ □= □139.4 (95% CI □= □19.637–445.564), *β*_*T8*_ □= □156.4 (95% CI □= □28.840–682.151), *β*_*T9*_ □= □58.8 (95% CI □= □11.709–290.651); Shannon: *β*_*T0*_ □= □ 5.2 (95% CI □= □ 3.642–5.844), *β*_*T1*_ □= □4.4 (95% CI □= □ 1.389–6.496), *β*_*T3*_ □= □3.3 (95% CI □= □0.654–5.011), *β*_*T4*_ □= □3.2 (95% CI □= □0.568–4.918), *β*_*T8*_ □= □3.0 (95% CI □= □0.776–5.306), *β*_*T9*_ □= □1.5 (95% CI □= □0.765–3.777)).

**Figure 3.**
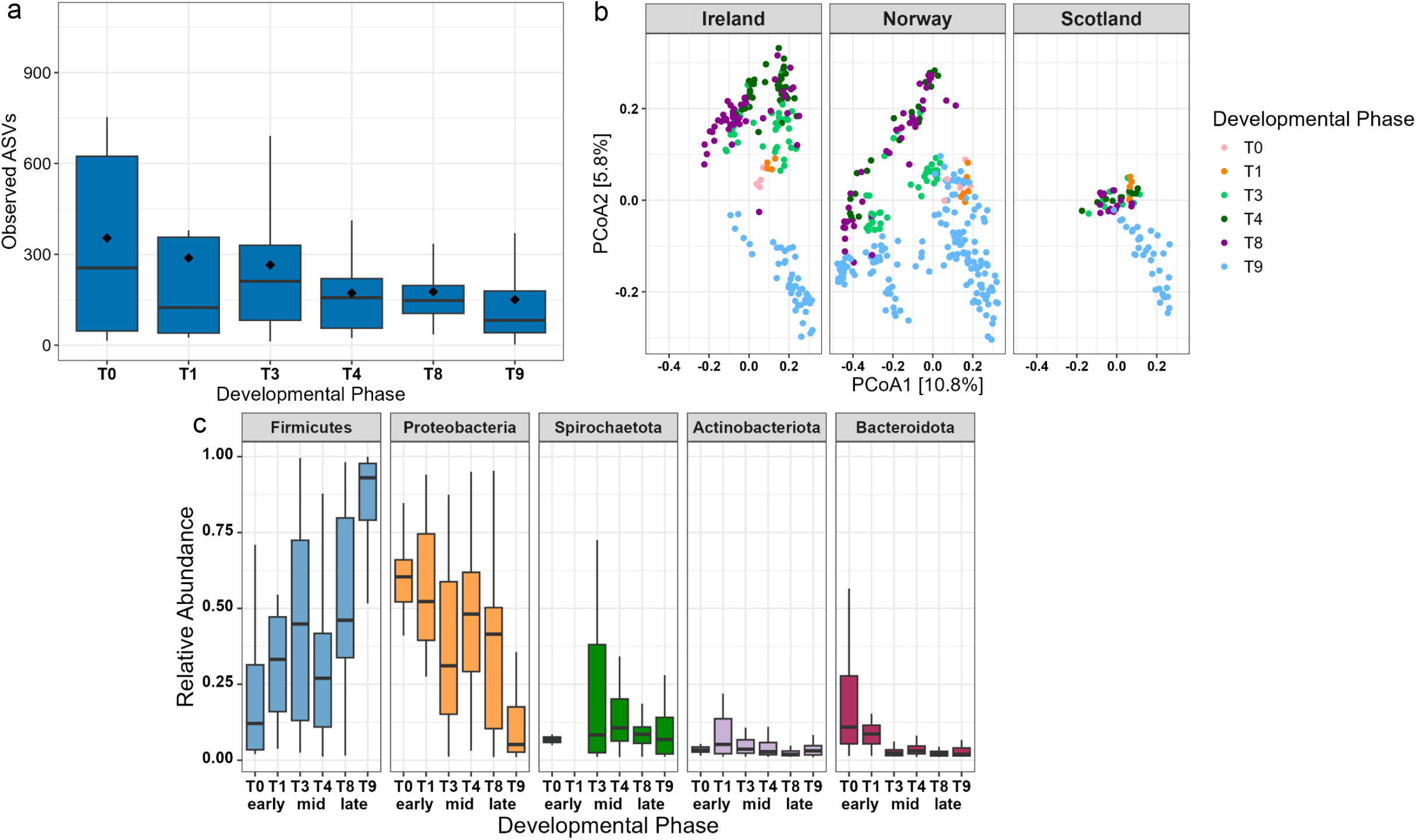
Farmed salmon microbiota differ in diversity and composition across developmental phases. (**a**) Box plots showing Observed ASVs for farmed salmon microbiota across developmental phases. The diamond, line symbols represent mean, median respectively. (**b**) The PCoA plot reflecting microbial community composition of farmed salmon microbiota across developmental phases. Percentages of explained variance by each principal axis are indicated in square brackets. (**c**) Box plot showing four major phyla (>0.1%), specifically, Firmicutes (ΔAIC_C_ = 42·68), Proteobacteria (ΔAIC_C_ = 31·06), Actinobacteriota (ΔAIC_C_ = 29·05) and Bacteroidota (ΔAIC_C_ = 39·02) differ in relative abundance across developmental phases.

Similar to alpha diversity, with PERMANOVA, we found that timepoint (R^2^ = □0.188, p = □0.001), but also country (R^2^ = □0.059, p = □0.001) and the interaction between them (country*timepoint; R^2^ = □0.037, p = □0.001), influenced the salmon microbial community composition. A similar direction of spread of data points for particular SDP across countries was observed in the PCoA plot (Figure 3b). Sequencing run (R^2^ = □0.110, p = □0.001) and sequencing depth (R^2^ = □0.003, p = □0.002) also significantly influenced the results in the PERMANOVA model.

### Differentially abundant major phyla and ASVs among salmon developmental phases

We assessed whether SDP could influence the relative abundance of taxa at phylum and ASV level. By running GLMMs at the higher phylum level (>0.1%), four phyla, Firmicutes (ΔAIC_C_ = □42.68), Proteobacteria (ΔAIC_C_ = □31.06), Actinobacteriota (ΔAIC_C_ = □29.05) and Bacteroidota (ΔAIC_C_ = □39.02), showed strong support for SDP. Specifically, we observed an increase in Firmicutes and a decrease in Proteobacteria abundance across successive SDPs (Figure 3c). The other two phyla, Actinobacteriota and Bacteroidota, showed varying abundance patterns across different SDPs.

Similar to phylum, we detected major linear trends of increasing, decreasing or mid-peak following the SDPs, i.e., early, mid and late. Seventy-three ASVs were found to be significantly (p≤0.05) different in abundance, in which the majority-41 decreased, 5 increased and 27 exhibited mid-peaks following successive SDPs (Figure 4). The decreasing ASVs belonged to the phyla Firmicutes (9 ASVs), Proteobacteria (21 ASVs), Actinobacteriota (6 ASVs), Bacteroidota (3 ASVs) and Planctomycetota (2 ASVs), whereas the increasing ASVs were all from Firmicutes (5 ASVs). Mid-peak ASVs were from Firmicutes (7 ASVs), Proteobacteria (11 ASVs), Actinobacteriota (8 ASVs) and Bacteroidota (1 ASV). One ASV belonging to *Mycoplasma* (ASV_061283) increased multiple fold (∼7000) from the early to late phase.

**Figure 4.**
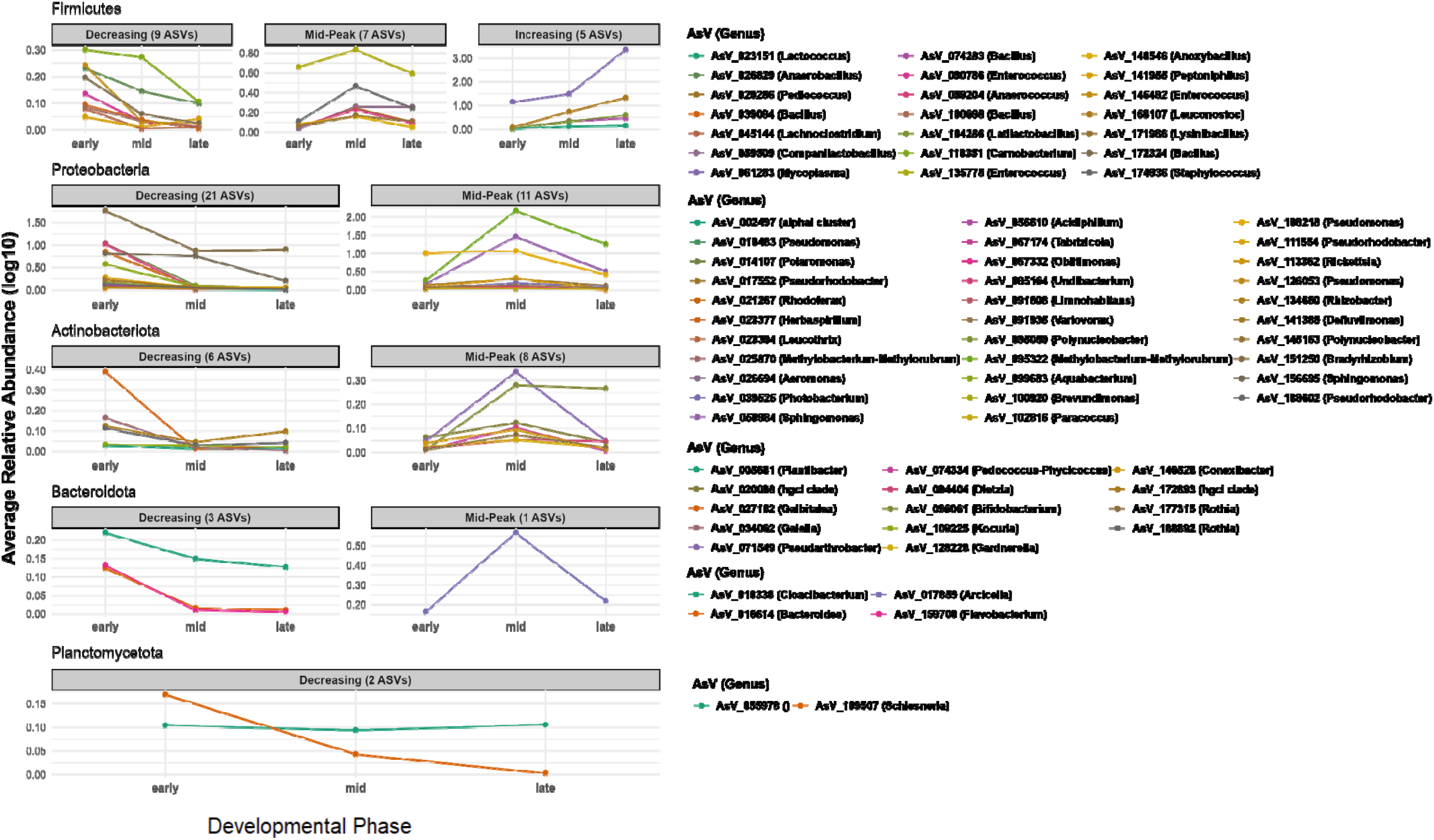
ASVs showing differentially abundant trends across developmental phases in farmed salmon gut microbiota. The line plots display the change in relative abundance of ASVs, across major developmental phases (early, mid and late) of farmed salmon on y- and x-axes, respectively, sorted according to their respective phylum with legend showing their genus identity. ASVs showing differential abundant trend (decreasing, increasing or mid-peak) in successive developmental phases were determined by ANCOM-BC2 (Holm-Bonferroni correction, p≤0.05).

### Diet, environment and farmed salmon gut microbiota diversity and composition

We compared microbial alpha diversity of diet, environment and farmed salmon gut (i.e., digesta, intestinal tissue) using a GLMM. We found strong support for an effect of country, sample category, and their interaction (country*sample category) on Observed ASVs, but not for timepoint (ΔAIC_C_ = □592.02, R^2^_GLMM(m)_ = □0.606, R^2^_GLMM(c)_ = □0.747, Figure 5a). Similar results were found using Shannon indices, except that a timepoint effect was also supported in the model (ΔAIC_C_ = □414.40, R^2^_GLMM(m)_ = □0.525, R^2^_GLMM(c)_ = □0.626). Microbial alpha diversity was highest for seawater samples, followed by freshwater, diet, digesta and intestinal tissue (Observed ASVs: *β*_*IT*_ □= □108.3 (95% CI □= □30.013–397.004), *β*_*DIG*_ □= □235.5 (95% CI □= □ 65.049–869.022), *β*_*DI*_ □= □488.8 (95% CI □= □246.209–972.53), *β*_*FW*_ □= □1918.0 (95% CI □= □510.781– 6814.801), *β*_*SW*_ □= □1972.1 (95% CI □= □503.968–7585.981); Shannon: *β*_*IT*_ □= □2.7 (95% CI □= □0.804– 4.728), *β*_*DIG*_ □= □3.9 (95% CI □= □1.965–5.911), *β*_*DI*_ □= □4.3 (95% CI □= □3.291–5.277), *β*_*FW*_ □= □5.9 (95% CI □= □3.899–7.851), *β*_*SW*_ □= □6.2 (95% CI □= □4.111–8.252)).

**Figure 5.**
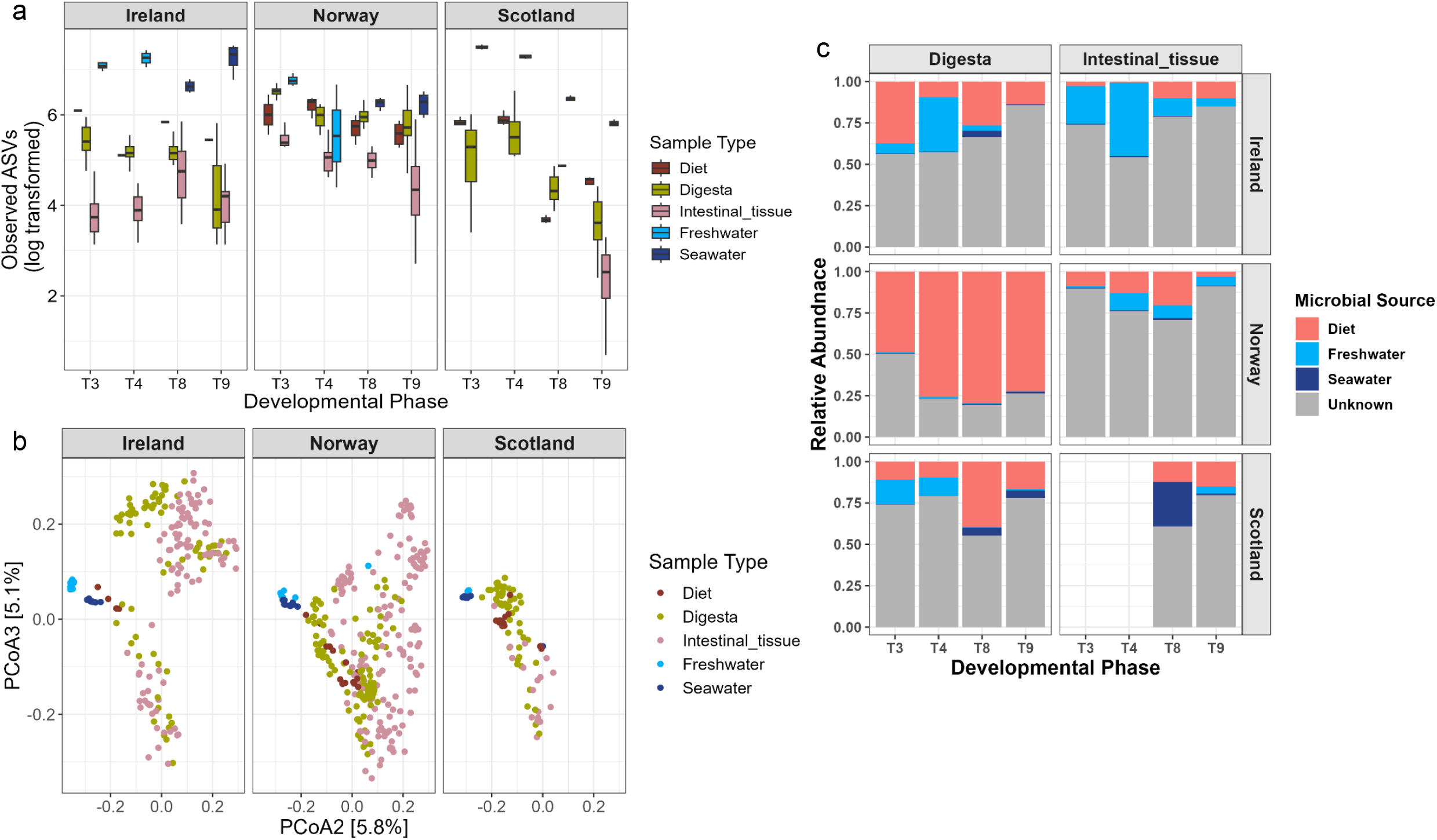
Microbiota of diet, environment and farmed salmon gut differ in diversity and composition across countries. (**a**) Box plots showing Observed ASVs (natural log transformed) in gut (digesta, intestinal tusse), diet and environment (freshwater, seawater) of farmed salmon across developmental phases and countries. (**b**) The PCoA plot reflecting microbial community composition gut (digesta, intestinal tusse), diet and environment (freshwater, seawater) of farmed salmon across countries. Percentages of explained variance by each principal axis are indicated in square brackets. (**c**) Bar plot showing relative contribution of diet and environment (freshwater, seawater) in the digesta and intestine tissue microbiota across developmental phases of farmed salmon across countries.

Similarly, with the PERMANOVA model, we found strong support for an effect of sample category (R^2^ = □0.156, p = □0.001), but also for country (R^2^ = □0.057, p = □0.001), and its interaction with timepoint (country*timepoint; R^2^ = □0.081, p = □0.001) (Figure 5b). Sequencing run (R^2^ = □0.207, p = □0.001) and sequencing depth (R^2^ = □0.002, p = □0.001) also significantly influenced the results in the model.

### Salmon gut microbiota shaped by diet and environment

To find out the potential source of salmon gut microbes through SourceTracker analysis, we observed that the majority of taxa are unique to either digesta or intestinal tissue (unknown=0.674 (95% CI □= □0.646–370.482)). In other words, for most of these microbes, the potential source of origin could not be identified. Overall, other than microbes of unknown origin, microbes from diet (0.237 (95% CI □= □0.210–0.264)) dominated the gut microbiota, followed by freshwater (0.079 (95% CI □= □0.064–0.095)) and seawater (0.009 (95% CI □= □0.005–0.013)) (Figure 5c). Regarding microbial source, we observed country (i.e. Ireland, Norway, Scotland), timepoint (i.e. T3, T4, T8, T9) and compartment (digesta or intestinal tissue) specific patterns. For example, in Norway, the digesta samples mostly contained microbes from the diet, while intestinal tissue samples contained microbes from both diet and environment (Figure 5c).

## Discussion

Understanding the gut microbiota of Atlantic salmon is essential for optimizing aquaculture practices and enhancing farmed fish health, as well as for better management and conservation of wild salmon populations^35,36^. This study provides a comprehensive analysis of the gut microbiota of wild and farmed Atlantic salmon, highlighting significant differences in microbial diversity, composition, and taxa abundances, influenced by factors such as biogeography, SDPs, and environmental exposure. As the gut microbiota plays a vital role in host health, immunity, and disease resilience, such information is crucial for both farmed fish in controlled aquaculture environments, as well as for wild populations.

### Wild salmon shows higher gut microbiota diversity and varying community composition in comparison to farmed salmon

Wild Atlantic salmon exhibited significantly higher gut microbial alpha diversity compared to farmed salmon, as indicated by the Observed ASVs and Shannon diversity indices, which differ according to salmon origin country (Norway, Ireland, and Scotland). The reduced microbial diversity in farmed salmon likely reflects the limited environmental conditions in aquaculture, where dietary and environmental microbial exposure is more restricted than in natural habitats. However, the gut microbiota could be further constrained by the limited genetic diversity of farmed salmon^37^. Similarly, higher gut microbial diversity in wild living individuals has been observed previously in diverse vertebrate species^38^, including Atlantic salmon^37^ compared to their captive or farmed counterparts. As habitat is an important driving factor in shaping gut microbial diversity, it is not surprising that country of origin influences microbiota diversity and that diversity differs according to country, with Norwegian salmon exhibiting the highest diversity. These differences most likely reflect local environmental microbiota that may colonize the fish gut (e.g., in wild) or vary based on country-specific aquaculture practices (e.g., for farmed), and emphasize the potentially significant role of regional factors in shaping microbial communities within the host^35,39^. Digesta samples exhibited higher microbial diversity than intestinal tissue samples, which is consistent with previous findings from mammals and fish^40,41^. This could be explained by host selection pressure on tissue-residing bacteria as well as colonization resistance by existing dominant bacteria colonizing intestinal tissues^42,43^.

Similar to the difference in alpha diversity, we identified that the overall microbial community composition differs between wild and farmed salmon, together with country and sample type. SDS (wild or farmed) can explain 8.9% of the variation alone, and 2% of the variation together with country and sample category (SDS*country*sample category), whereas country can explain 7.1% and sample category 5.3% of the variation in beta diversity models. This suggests that SDS, being from wild or farm origin, can give rise to differences in their microbial community composition, which is more than the effect of country or sample category on microbial community. The more controlled conditions in aquaculture, including diet and water treatments (particularly during the freshwater phase), may give rise to microbial communities different from those in the natural environments, as observed previously^37^. Effects of geography, habitat and sample category on microbial community composition have been previously reported in diverse fish species^35,39^. Questions derived from these observations are: Which bacterial taxa differ between wild and farmed salmon across countries? And is there any commonality in such patterns across countries?

Differences in relative abundance were observed at the phylum level, with Firmicutes and Bacteroidota being more abundant in farmed salmon and Proteobacteria showing higher abundance in wild fish. This shift is consistent with previous findings suggesting that Firmicutes often increase under controlled diets, reflecting an adaptation to the higher carbohydrate content common in aquaculture feeds^16^. Conversely, wild salmon’s higher Proteobacteria levels may indicate a broader environmental exposure that supports a wider range of microbial taxa^39^. At the ASV level, several taxa, including *Mycoplasma, Aliivibrio*, and *Photobacterium*, were more abundant in farmed salmon across all three countries, whereas *Brevinema* and *Rickettsia* were consistently lower. *Mycoplasma, Aliivibrio, Photobacterium*, and *Brevinema* are often dominant commensals in salmon, and their differential abundance could reflect specific adaptations to aquaculture diets and environments, aligning with prior findings of microbiota shifts in response to diet and environmental containment^35,44,45^.

### Influence of the salmon developmental phases on the gut microbiota dynamics in farmed salmon

Investigating microbiota across farmed salmon SDPs, we observed a consistent decrease in microbial diversity (i.e. Observed ASVs, Shannon) corresponding to increasing age, which is further influenced by salmon’s country of origin. Such age-associated reduction in microbial diversity may be attributed to immune system maturation, which increases host-driven selection pressures on the microbiota, as well as dietary shifts across SDPs and salinity-related physiological changes during the transition from freshwater to seawater^35^. Immune system can actively regulate microbial diversity or indirectly favour the proliferation of specific taxa, such as *Mycoplasma*, leading to an overall reduction in diversity^17^. Differences in farming practices across countries such as diet, treatments further influenced microbiota diversity at specific timepoints; however, the overall trend of age-related decline in diversity remained consistent across all studied countries. One limitation of our sampling is that, for the two earliest SDPs: fertilized eggs (T0) and yolk sac larvae (T1), DNA was extracted from pooled samples comprising 25 eggs or yolk sac larvae per sample. This approach was necessary due to the low amount of starting material and the potentially low microbial load associated with individual eggs, likely a result of routine surface disinfection upon hatchery introduction. The pooling may have increased the observed within-sample diversity compared to the actual diversity present within individual eggs/larvae. Nevertheless, a decrease in diversity continued in successive SDPs where individual fish were sampled (i.e., T3, T4, T8, T9), across countries. We observed a decrease in microbial diversity in both the diet and environment across successive SDPs of farmed salmon. This suggests that the observed decline in gut microbial diversity in farmed salmon is driven by multiple contributing factors.

Both SDP and farming environment influenced gut microbial community composition, with similar patterns emerging across countries for successive life phases. These results reinforce that both intrinsic factors (e.g., SDP) and extrinsic factors (e.g., environment) significantly shape gut microbiota composition in farmed salmon^46^.

Changes in microbial diversity and composition across SDPs were largely driven by significant shifts in the abundance of four major phyla: Firmicutes, Proteobacteria, Actinobacteriota, and Bacteroidota. Specifically, Firmicutes exhibited an increasing trend, while Proteobacteria displayed a decreasing trend in abundance across successive SDPs. These observed specific changes may be attributed to shifts in diet and environment across SDPs; for instance, the increase in Firmicutes likely reflects an adaptation to higher carbohydrate contents typical of aquaculture feeds^16^. A similar contrasting pattern between Firmicutes and Proteobacteria during salmon SDPs has been reported previously^47^. However, whether this pattern is associated with immune system maturation remains unclear and requires further investigation. For example, at the post-vaccination stage (T4), we observed a slight deviation in the abundance trends for both Firmicutes and Proteobacteria, suggesting that the vaccination regimen may influence these specific phyla and implicate a potential role of the immune system. This observation warrants further investigation in future studies. At the ASV level, the majority (∼56%) of ASVs exhibiting significant differential abundance trends showed a decrease in abundance across successive SDPs, consistent with our observation of reduced microbial diversity with advancing SDPs. Most of the decreasing ASVs belonged to Proteobacteria, whereas the increasing ASVs were associated with Firmicutes, consistent with the overall phylum-level trends. Notably, one Firmicutes ASV, specifically assigned to *Mycoplasma*, displayed the highest increase (∼7000 fold) in abundance, which may contribute to the observed reduction in overall gut microbial diversity, as discussed above^17^. *Mycoplasma* spp. are known to colonize the Atlantic salmon gut at different SDPs, and co-diversification with its salmonid host has been reported^48,49^. Mid-peak ASVs (representing 36% of all significant ASVs) were primarily affiliated with Firmicutes (7 ASVs), Actinobacteriota (8 ASVs), and Proteobacteria (11 ASVs). These ASVs may be influenced by factors such as diet, vaccination, or physiological transformations occurring during the mid-developmental phase of salmon, and further research is needed to clarify these associations.

### Environment and diet together shape the farmed salmon gut microbiota

Comparison of alpha diversity across various sources revealed that environmental samples (e.g., seawater, freshwater) exhibited the highest diversity, followed by diet, digesta, and intestinal tissue samples. These findings indicate that environmental sources provide a more diverse microbial pool compared to those introduced to the gut through diet, where selective pressures likely favour specific microbial taxa adapted to the gut environment^46^. Microbial community composition varied according to sample category, with diet samples appearing closer to the salmon gut microbiota compared to the environmental microbiota (Figure 5b). The key question derived from these observations is: To what extent is the gut microbiota (digesta or intestinal tissue) shaped by diet and environment?

We found that while both environment- and diet-associated taxa contribute to shaping the gut microbiota of Atlantic salmon, diet exerts a predominant influence over the environment. This reliance on diet-associated microbes underscores the strong impact of controlled aquaculture feeding practices on the microbiota and highlights the importance of carefully formulated diets to support beneficial microbial communities^50^. Nevertheless, the potential source of origin for the majority of gut microbes (67%) could not be identified, suggesting that these microbes are salmon gut-specific and that their sources were not included in our study. For example, we included a limited number of diet and environmental samples collected at specific SDPs. The contribution of microbial taxa differed by the compartment (digesta or intestinal tissue), SDP under consideration, and country of origin. For instance, in Norway, digesta samples predominantly contained bacteria derived from diet, irrespective of SDP. In contrast, in Scotland, digesta samples showed an equal contribution of bacteria from freshwater and diet at T3 and T4 stages, whereas at T8 and T9 stages, bacteria were primarily derived from diet and seawater. Due to a lack of intestinal tissue samples at T3, T4 stages for Scotland, we could not estimate the relative contribution of diet and environment taxa for these samples. These differences in microbial source patterns across countries and SDPs suggest that tailored aquaculture practices may be necessary to optimize microbiota health in varying contexts.

## Conclusion and Future Directions

This study provides a comprehensive view of the ecological dynamics of the Atlantic salmon gut microbiota. Our findings suggest that the lower microbial diversity observed in the gut of farmed salmon reflects a more specialized microbiota adapted to the aquaculture environment, which may have implications for fish health as microbiota diversity is often associated with a robust immune system. The SDP-specific microbial taxa patterns in farmed salmon, including the proliferation of *Mycoplasma* linked to reduced diversity, underscore gaps in our understanding of the origins and functions of salmon gut microbes, presenting opportunities for further research. Since diet is the predominant contributor to the gut microbiota, tailored strategies, such as region-specific diets, alternative feeding practices, and environmental management, can help maintain microbiota diversity and potentially enhance fish health. Future research should focus on the functional roles of the specific taxa identified here, particularly those differing consistently between wild and farmed salmon, to better understand their contributions to host health and resilience. Additionally, incorporating functional metagenomics and expanding microbiota studies beyond the gut to other body surfaces (e.g., skin and gills) could provide deeper insights into how these microbial communities influence host metabolism and immunity, ultimately guiding best practices for sustainable aquaculture.

## Author contributions

PL, SW, SM, AHJ acquired funding and resources. AHJ coordinated the Salmon research activities in the CIRCLES project. SG, SW, SM, OB provided host data. PL, FDM carried out 16S rDNA sequencing. HPK, W performed bioinformatics. W validated data, performed statistical analysis and wrote the manuscript. All authors commented and approved the submitted version.

## Data availability

The individual gut bacterial 16S rRNA gene sequences are available under European Nucleotide Archive under project ID PRJEB81272.

## Acknowledgements

This study was funded by the European Commission through the Horizon 2020 grant agreement no. 818290 (project CIRCLES) and by the Research Council of Norway. We thank Nordlaks smolt AS, Hamarøy, Norway (CIRCLES partner), for providing access to their production facilities for sampling of farmed salmon, and the Norwegian Environment Agency for permissions to sample wild salmon in Norway. We thank Lisa Furnesvik and Siw Larsen (NVI Harstad), Siri Kristine Gåsnes, Tine Tønder and Vegard Sollien (NVI Trondheim) for help with sampling, and Elin Johanne Trettenes and Kristin Soetaert (NVI Ås) for help with DNA extraction, Hanne Nørgaard Nielsen (DTU) for help with sequencing, Benjamin G. Clokie, E. McDonald and A. Elsheshtawy (UoS) for sampling, sample selection, processing and shipment. We thank the project partners and in particular all WP5/6 partners for valuable discussions.

## Ethical permission

All samples from Scotland were collected in accordance with the United Kingdom Scientific Procedures Act and approved by the University of Stirling Animal Welfare and Ethical Review Body (AWERB 2020 0270 153), wild salmon samples were collected with the permission of The River Deveron District Salmon Fishery Board. The Irish studies were carried out under the Health Products Regulatory Authority (HPRA) licence number AE19130-P056. In Norway the wild fish were captured at river Vigda in Skaun, Trøndelag as part of a spawning fish count by Norwegian Institute for Nature Research and local people and the fish sampling permission were obtained from Trøndelag county governor (Fylkesmannen), and at river Kjerringnes in Sortland, Nordland with the permission of Morten Halvorsen, State Administrator Nordland (Statsforvalteren), after clarification with Tore Vatne at the State Administration and the landowners’ association.

