## Supplement for "Ecological dynamics of the Atlantic salmon gut microbiota across developmental phases and geographic regions"

### Slide 1
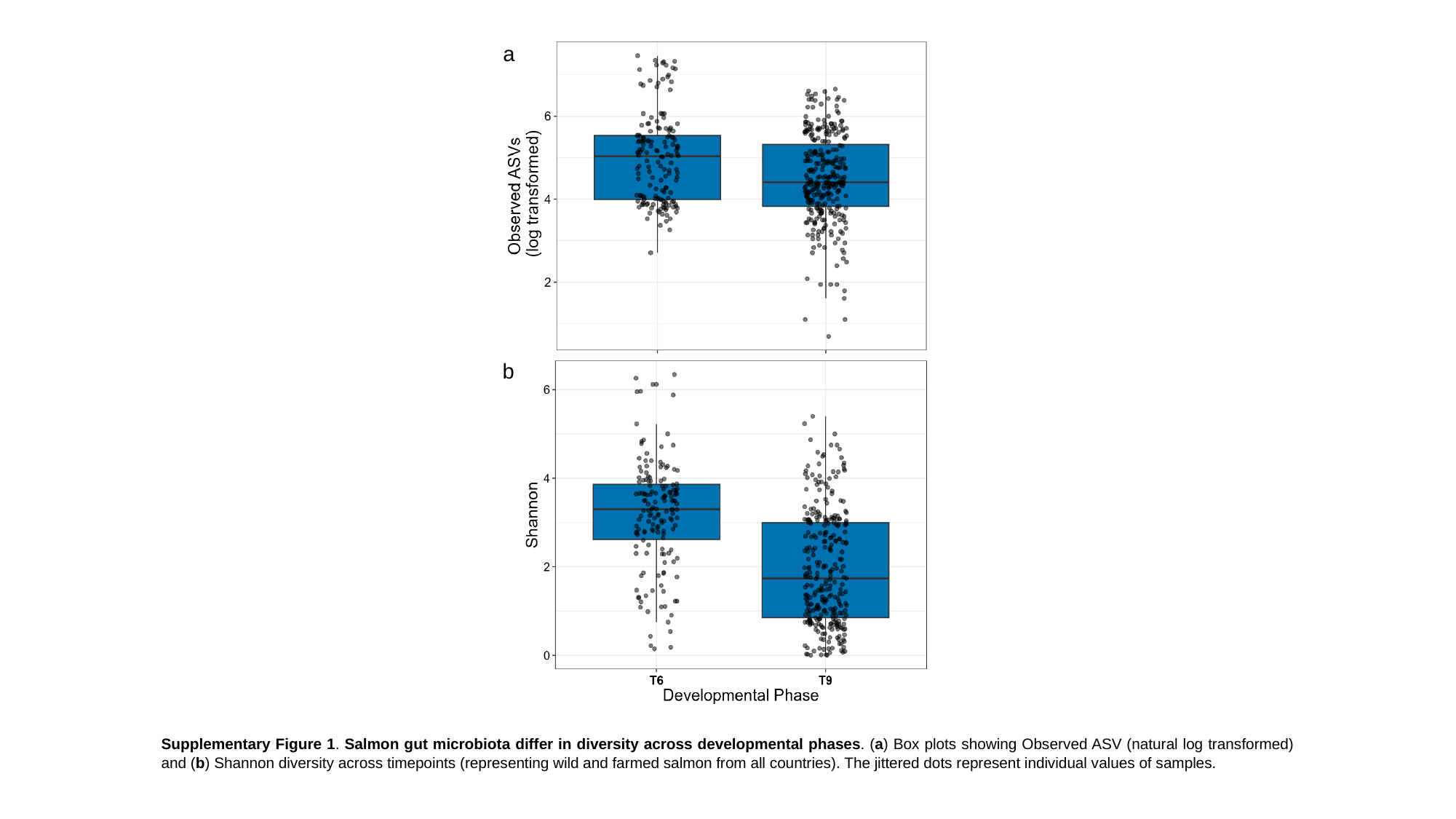

a
b
Supplementary Figure 1. Salmon gut microbiota differ in diversity across developmental phases. (a) Box plots showing Observed ASV (natural log transformed) and (b) Shannon diversity across timepoints (representing wild and farmed salmon from all countries). The jittered dots represent individual values of samples.

### Slide 2
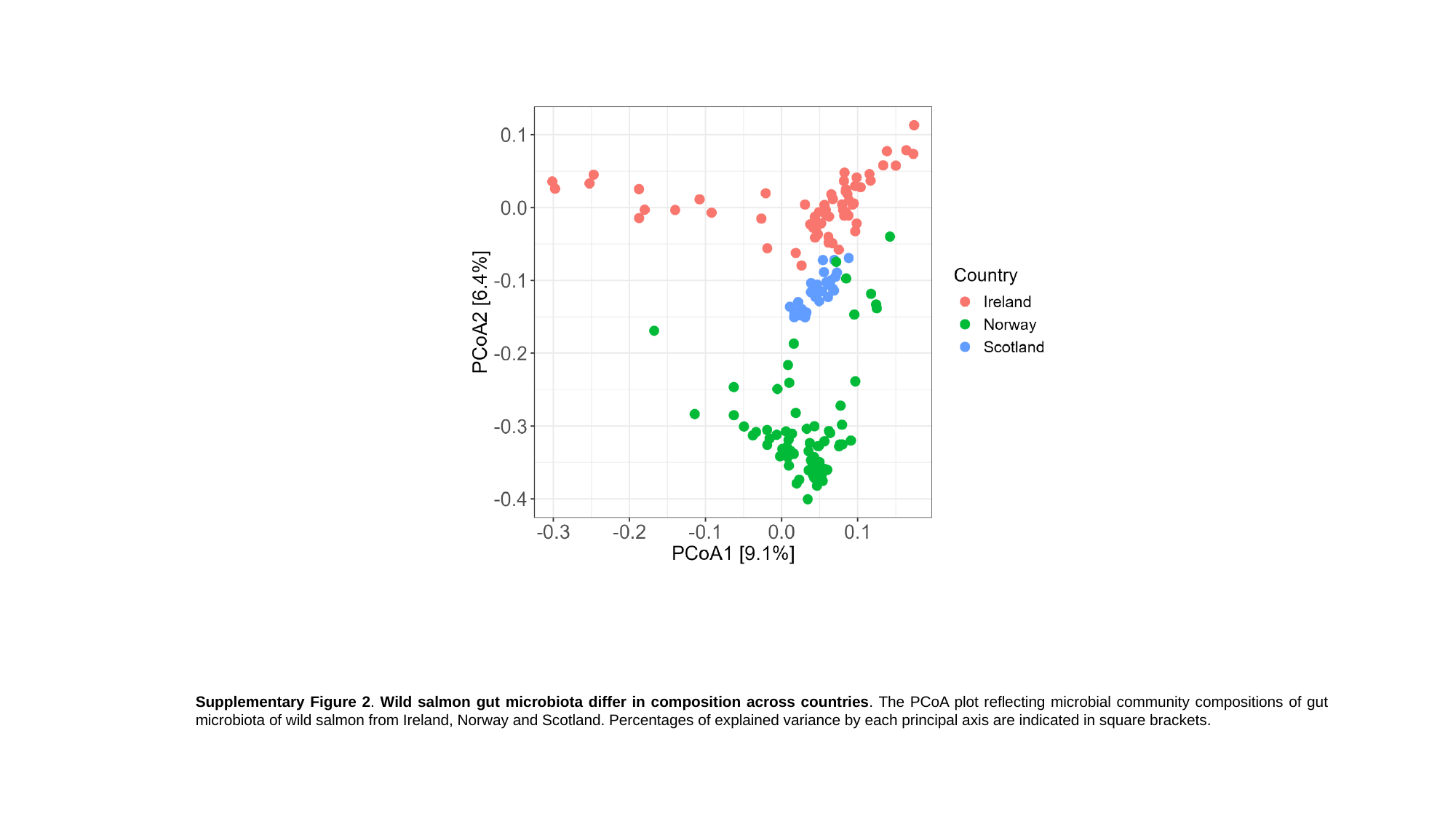

Supplementary Figure 2. Wild salmon gut microbiota differ in composition across countries. The PCoA plot reflecting microbial community compositions of gut microbiota of wild salmon from Ireland, Norway and Scotland. Percentages of explained variance by each principal axis are indicated in square brackets.

### Slide 3
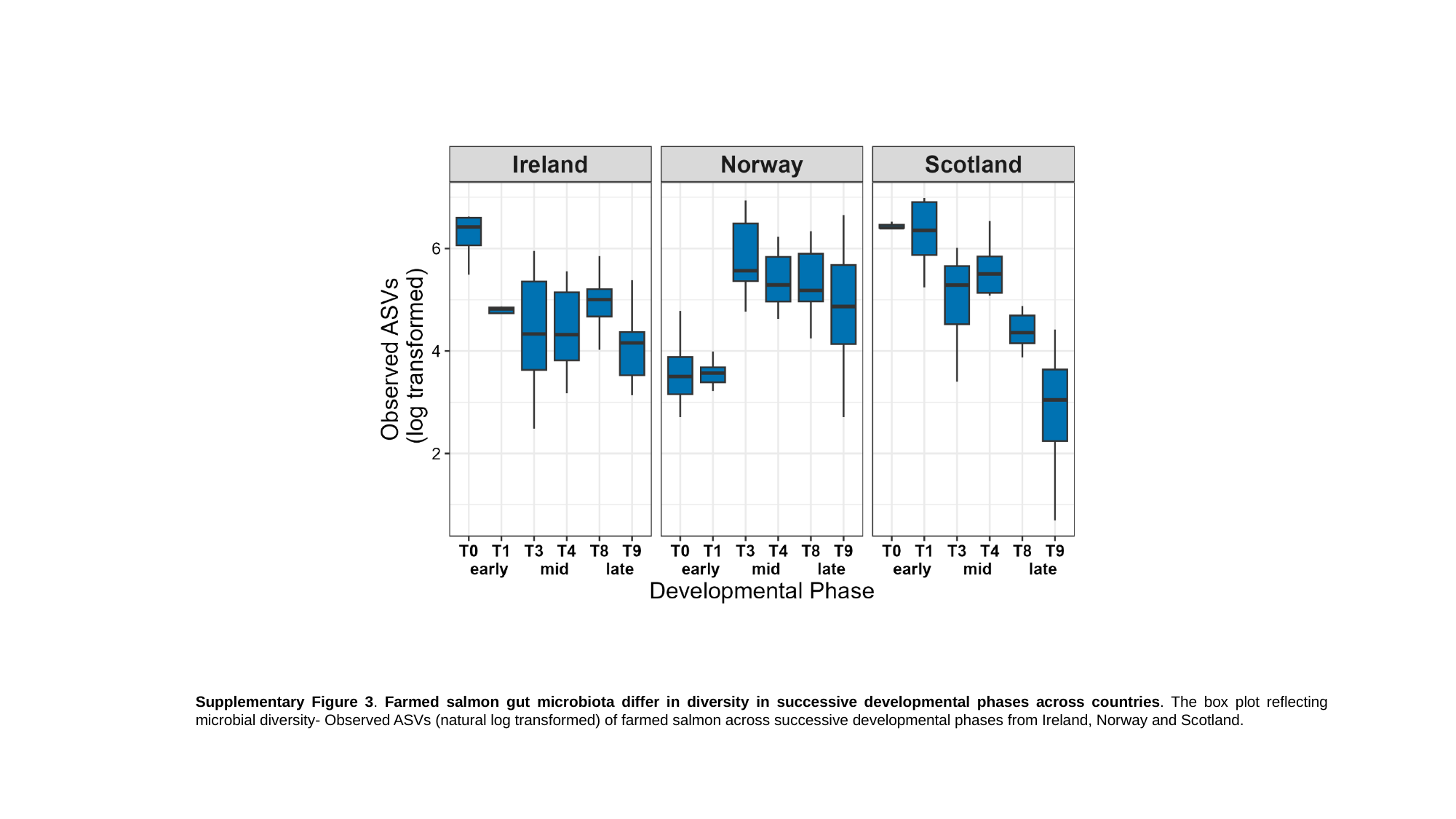

Supplementary Figure 3. Farmed salmon gut microbiota differ in diversity in successive developmental phases across countries. The box plot reflecting microbial diversity- Observed ASVs (natural log transformed) of farmed salmon across successive developmental phases from Ireland, Norway and Scotland.
